# Fractionated ionising radiation affects cellular functions, and gene expression associated to subpopulation of F11 dorsal root ganglia neurons without inducing oxidative stress

**DOI:** 10.64898/2026.08.11.735255

**Authors:** William Timbury, Sean M. Gettings, Raymond Shek, Craig D. Lindsay, Riddhi Sharma, Mustafa Najim, Nora Bourbia

**Affiliations:** UK Health Security Agency, Radiation Effects Department, Radiation Protection Science Division, Harwell Science Campus, Didcot, Oxfordshire OX11 0RQ, UK; Johnston Cancer Research Centre (JCRC), Queen’s University Belfast, Belfast, BT9 7AE, UK

**Author notes:** Corresponding author: Nora Bourbia.

**Keywords:** nociception, ionising radiation, radiotherapy, dorsal root ganglion, sensory neuron, X-ray

## Abstract

Radiotherapy is common practice to treat cancer but produces significant side effects such as chronic pain. Cancer survivors report developing chronic pain due to their treatment even long after the cancer is cured. To understand the mechanisms underlying the radiotherapy-induced chronic pain, we assessed how ionising X-ray radiation exposure during 4 consecutive days of 5 Gy (total radiation dose of 20 Gy) affected dorsal root ganglia (DRG) sensory neurons (rodent F11 cell line). On the 5^th^ day, we assessed known impacts of ionising radiation (senescence, oxidative stress, cellular metabolism, mitochondrial copy number, and mitochondrial respiration) followed by assessing expression of genes associated with populations of DRG neuronal fibres. We discovered that fractionated exposure to ionising radiation increased senescence, mitochondrial copy number, and modulated the NAD^+^/NADH pathway, but did not change the oxygen consumption rate nor induce oxidative stress 24 hours after the last irradiation exposure. Additionally, ionising radiation altered the expression of genes associated with mechanoreceptor fibres, known to have pro-nociceptive properties in the context of injury and chronic pain.

## 1. Introduction

Radiotherapy is a common cancer treatment using ionising radiation to kill cancer cells however it also produces significant side effects such as chronic pain (Karri *et al*., 2021). Indeed, 33% of cancer survivors report chronic pain (i.e.: myelopathy, fibrosis, peripheral nerve entrapment, plexopathy) due to cancer treatments including radiation therapy (Polomano & Farrar, 2006; Paice, 2011; Pachman *et al*., 2012; Glare *et al*., 2014). A recent overview of radiotherapy-induced chronic pain in childhood cancer survivors showed that radiotherapy can cause neuropathy by radiation damage to the nerves or surrounding tissues, damaging the nerves (Chua & Vig, 2023). Current medications are moderately effective at best with extensive side effects (Cavalli *et al*., 2019). Chronic pain is also associated with a range of comorbidities, including depression (Bourbia & Pertovaara, 2011; Sheng *et al*., 2017), negatively impacting the lives and wellbeing of children and adolescents (Koechlin *et al*., 2020; Chua & Vig, 2023). Therefore, it is important to study how radiotherapy induces chronic pain to prevent such complications.

In this study we exposed the F11 cell line (hybrid rodent cell line between rat dorsal root ganglion (DRG) neurons and mouse neuroblastoma cells) to either sham-irradiation or X-ray irradiation at a dose of 5 Gy per day for four consecutive days, resulting in a cumulative dose of 20 Gy. On the fifth day, we evaluated whether various cellular processes were influenced by radiation (senescence, mitochondrial copy number, mitochondrial respiration, NAD+/NADH ratio, and oxidative stress). To determine whether irradiation affected specific characteristics of DRG neurons, we further assessed the expression of genes associated with sensory fibres across the 17 DRG neuron subpopulations described by Jung and colleagues (Jung *et al*., 2023)

## 2. Materials & methods

### 2.1. Cell culture

Undifferentiated F11 (cat: 08062601-1VL) were cultured in either T25 flasks, 6-well plates (seeding density of 200,000 cells/well), or 96-well plate (seeding density of 20,000 cells/well) using a cell culture medium composed of DMEM high-glucose (cat: D6429-6X500ML; Merck) complemented with 2 mM L-Glutamine (cat: 25030-024; supplier: ThermoFisher Scientific), 10% FBS (cat: 10500064; Thermofisher Scientific), and 1% penicillin-streptomycin (cat: P4333-100ML; Merck). Cells were incubated at 37⁰C and 5% CO_2_.

### 2.2. X-ray irradiation

Cells were exposed to either sham-irradiation or a daily single dose of 5 Gy X-ray radiation (at a dose rate of 1.7 Gy per minute for 2.9 minutes) for four consecutive days using an X-ray source from AGO X-ray Limited controlled by AGO HS MP1 X-ray controller. On the fifth day, cells were either pelleted and stored in -80°C or processed for further assays. In a separate experiment, cells were exposed to one single dose of 5 Gy or sham exposure, and the cells were pelleted 5 hours later for further assay.

### 2.3. Oxygen consumption rate (OCR)

Oxygen consumption rate (fmol/mm²/s) was assessed using the Resipher 32-channel equipment by Lucid Scientific. Microplate strips (cat: 11653189, supplier: BRAND^TM^) were used to allow the recording of both treatments (sham and X-ray-irradiated) on the same 96-well plate with the Resipher. Treatments were randomly allocated among the 4 strips used (2 X-ray and 2 sham).

OCR was assessed 14h after the sham-or X-ray-radiation exposure by averaging a 3-hour period window. 14h after the exposure represented a stable recording of OCR. Cell confluency (%) was recorded before the 3^rd^ and 4^th^ sham radiation exposure, as well as 24h after the 4^th^ exposure using Incucyte® live-cell analysis system. Confluency was used to normalise the OCR (n = 8 to 16 per experimental group). No confluency was measured at prior time points. 2 strips (1 X-ray and 1 sham strip) were out of focus during the live-cell imaging scan which assessed cell confluence before the 3^rd^ exposure. Therefore, these were both excluded from analysis at this timepoint.

### 2.4. Mitochondrial copy number

Relative mitochondrial copy number was assessed using the quantitative polymerase chain reaction (qPCR) method as previously described (Gettings *et al*., 2024). DNA was extracted from the cells using the DNeasy Blood & Tissue Kit (cat.: 69504; Qiagen). 50 ng of DNA was then used to perform a qPCR using PerfeCTa SYBR Green SuperMix (cat.: 733–1246; VWR International) for an initial 95°C for 10 min followed by 40 cycles (95°C for 15 s, followed by 62°C for 60 s, and then 72°C for 20 s) in a Magnetic Induction Cycler (Mic qPCR, Bio Molecular Systems).

Two primer sets (Guo *et al*., 2009) were used to assess relative mitochondrial copy number and were tested in prior experimentation to validate their efficiency: the NADH:Ubiquinone Oxidoreductase Core Subunit V1 (*Ndufv1)* for the nuclear DNA (Forward sequence: 5’ -CGGGTATCTGTGCGTTTCAG -3’ and reverse sequence: 5’ -GTGTCTTGTACCAGTCACCCC -3’), and cytochrome c oxidase subunit I (*Mt-co1)* for the mitochondrial DNA (Forward sequence: 5’ -CTAGCCGGAAATCTAGCCCA -3’ and reverse sequence: 5’ -TGCGGCTAGTACTGGTAGTG -3’).

The relative mitochondrial DNA content was determined using the 2 x 2^ΔCT^ method (ΔCT = nuclear DNA CT – mitochondrial DNA CT) (Rooney *et al*., 2015). N = 6 per experimental group.

### 2.5. Senescence

Senescence was measured using the β-galactosidase assay (cat: 9860S; Cell Signalling Technology) combined with Fluorescein Di-β-D-Galactopyranoside (FDG) (cat: F2756-25MG; Sigma-Aldrich). Briefly, cells in 12-well plates were washed and trypsinised (cat: T4174-100ML; supplier: Merck) followed by inhibition with soya-bean trypsin inhibitor (cat: R007100; Thermofisher Scientific). The cell suspensions were then spun down and resuspended with reaction buffer composed of: 10x staining buffer, 100x solution A, 100x solution B provided within the β galactosidase kit, and with 200 mM of FDG and 10X Triton X-100 (cat: T8787-250ML; supplier: Merck). The reaction buffer was topped up with sterile water (cat: W1509-500ML; supplier: Merck) to reach a volume of 200 µL per well. After 1 hour incubation at 37⁰C, fluorescence was measured using a plate reader with excitation at 485 nm and emission at 528 nm. Exposure to 10 µM etoposide during the first three days, served as the relevant positive control. Cells were counted using a haemocytometer (cat: DHC-N01; supplier: Scientific Laboratory Supplies) to normalise the fluorescence per cell. N = 5 per experimental group.

### 2.6. Protein extraction and quantification

Cell pellets were processed for protein extraction using 350 µL of Pierce lysis buffer (cat. 87787, supplier: ThermoFisher Scientific) with 3.5 µL of Halt Protease Inhibitor Cocktail (cat. 78430, supplier: ThermoFisher Scientific). After 5 minutes at room temperature, the samples were centrifuged 13,000g at 4°C for 10 minutes. The supernatants were then transferred into a QIAshredder column (cat. 79656, supplier: QIAGEN) and centrifuged for 1 minute at 13,000g to yield the protein extract. The Pierce BCA Protein Assay Kit (Cat. A65453, supplier: ThermoFisher Scientific) was used to quantify the concentration of protein in a 20 µl aliquot of the extract. The manufacturer protocol was followed using 20 µl of samples or albumin standards. Albumin standards were 0 (blank), 50, 70, 100, 300, 500, 700, and 1000 µg/mL. Samples and standards were transferred in triplicates into a 96-well plate for absorbance (562 nm) readings.

### 2.7. Superoxide dismutase (SOD) activity

SOD assay kit (cat: MAK528, Sigma-Aldrich) was used to assess whether the SOD-dependant cellular antioxidative system (Landis & Tower, 2005) is impaired. The cell pellets were washed with HBSS (cat. 14025092, supplier: ThermoFisher Scientific) and resuspended in Pierce lysis buffer. Then the experiment followed manufacturer protocol. The SOD enzyme activity is measured in U/mL and then normalised to the protein concentration (µg/mL). N = 5-6 per experimental group.

### 2.8. Carbonyl groups quantification

Carbonyl assay kit (cat: MAK486, Sigma-Aldrich) was used to quantify carbonyl groups, which indicates the level of oxidative posttranslational modification of proteins, primarily driven by reactive oxygen species. The experiment followed manufacturer protocol using 100 µL of protein. The carbonyl content was normalised to the protein concentration to measure carbonyl protein (nmol/mg). N = 6 per experimental group.

### 2.9. NAD^+^/NADH Assay

NAD^+^/NADH Assay Kit (cat: MAK468, Sigma-Aldrich) was used to assess the concentration of NAD^+^ and NADH normalised to the cell confluency measured prior the assay, and to determine the ratio of NAD^+^/NADH. The manufacturer protocol was followed with the use of PBS (cat. P4417-100TAB, Sigma-Aldrich) diluted in MilliQ water, and nuclease-free water (Cat. AM9937, Supplier: Thermofisher Scientific). N = 6 per experimental group.

### 2.10. RNA extraction and quantification

The RiboPure RNA Purification Kit (cat. AM1924, supplier: ThermoFisher Scientific) was used to extract and purify the RNA. Briefly, RNA was extracted from the cell pellet by homogenising the cells using TriReagent (cat. AM9738. Supplier: ThermoFisher Scientific) followed by the addition of 1-Bromo-3-Chloropropane (Cat. B9673-200ML supplier: Merck). Downstream processing was done according to the manufacturer’s protocol. Quantification of RNA was done using 1 µL of extracted RNA on the NanoDrop Lite spectrophotometer by Thermo Scientific.

### 2.11. cDNA conversion

2,000 ng of RNA was converted in cDNA following the manufacturer protocol (without RNAse inhibitor) of High-Capacity cDNA Reverse Transcription Kit (Cat. 4368814, supplier: ThermoFisher), with a thermocycling protocol of 25°C for 10 min, then 37°C for 120 min, then 85°C for 5 min. The cDNA was stored in -20°C until further processing.

### 2.12. Gene expression using reverse transcription quantitative polymerase chain reaction (RT-qPCR)

PerfeCTa SYBR Green SuperMix (cat. 733-1250, supplier: VWR) with primer sets (Table 1) was used to assess gene expression between the experimental groups. The manufacturer’s protocol was followed by mixing 10 µL of PerfeCTa SYBR Green SuperMix with 1 µL of each forward and reverse primers (primers at a concentration of 10 µM), 3 µL of nuclease-free water (Cat. AM9937, Supplier: Thermofisher Scientific), and 5 µL of cDNA. RT-qPCR was performed in a Magnetic Induction Cycler (Mic qPCR, Bio Molecular Systems) set to an initial 90°C for 2 min followed by 40 cycles (95°C for 5 s, followed by 58°C for 15 s, and then 72°C for 10 s). *Hprt1* was used as housekeeping gene to determine the delta cycle threshold (ΔCT). Statistical analysis was performed on the ΔCT to compare both experimental groups. Following a normality test using Shapiro-Wilk, unpaired t-tests or Mann-Whitney tests were used to compare experimental groups (n = 5 to 6 per experimental group). Relative quantification (RQ) was used for graphical representation of the fold change between the experimental groups.

**Table 1:**
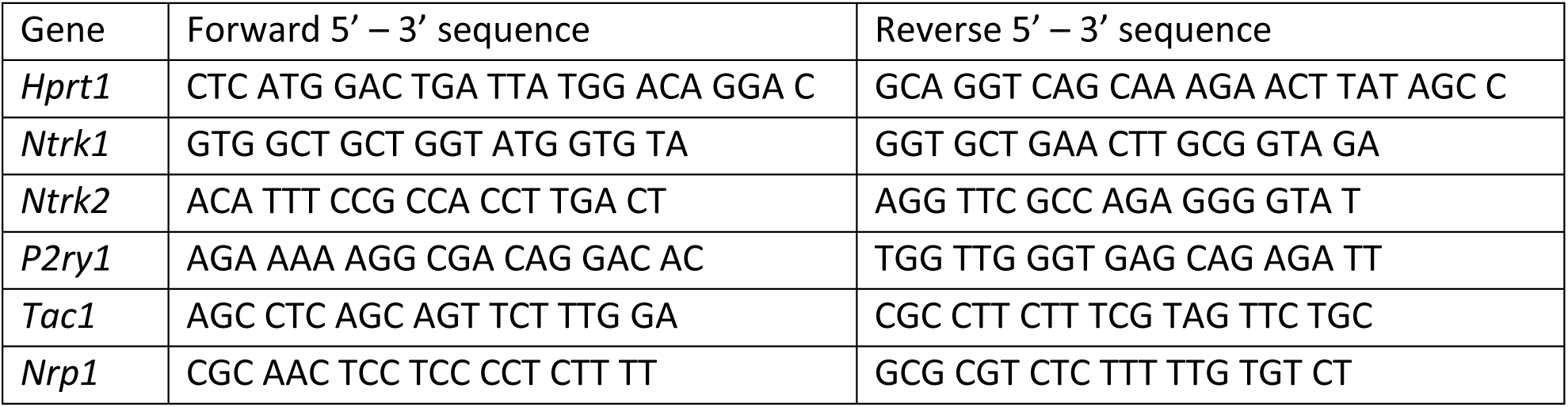
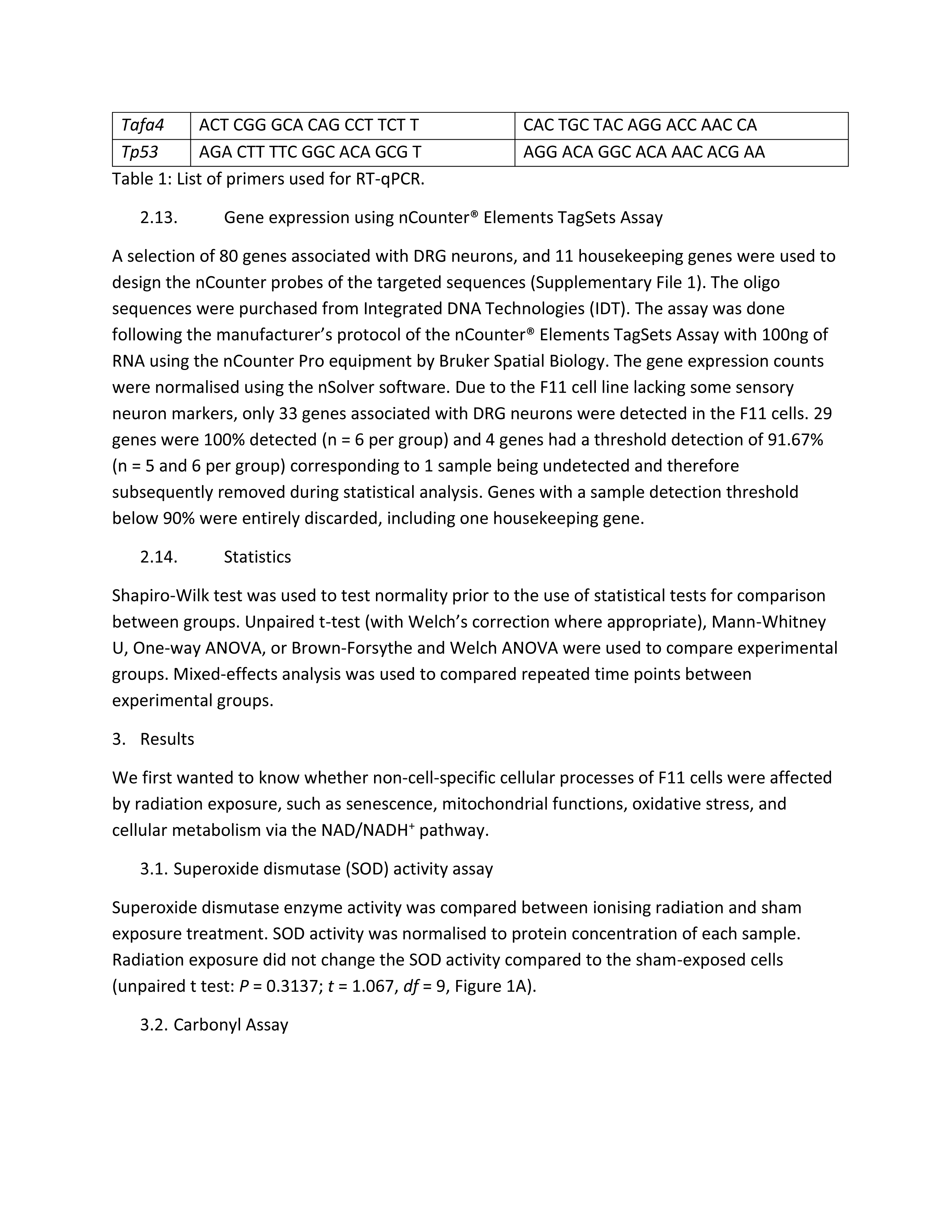
List of primers used for RT-qPCR.

F11 is a rat embryonic dorsal root ganglion (DRG) and mouse neuroblastoma hybrid cell line, therefore the primers have been designed using the webtool mrprimerw2.com selecting the species *rattus norvegicus*. The primers were then validated to confirm their efficiency.

### 2.13. Gene expression using nCounter® Elements TagSets Assay

A selection of 80 genes associated with DRG neurons, and 11 housekeeping genes were used to design the nCounter probes of the targeted sequences (Supplementary File 1). The oligo sequences were purchased from Integrated DNA Technologies (IDT). The assay was done following the manufacturer’s protocol of the nCounter® Elements TagSets Assay with 100ng of RNA using the nCounter Pro equipment by Bruker Spatial Biology. The gene expression counts were normalised using the nSolver software. Due to the F11 cell line lacking some sensory neuron markers, only 33 genes associated with DRG neurons were detected in the F11 cells. 29 genes were 100% detected (n = 6 per group) and 4 genes had a threshold detection of 91.67% (n = 5 and 6 per group) corresponding to 1 sample being undetected and therefore subsequently removed during statistical analysis. Genes with a sample detection threshold below 90% were entirely discarded, including one housekeeping gene.

### 2.14. Statistics

Shapiro-Wilk test was used to test normality prior to the use of statistical tests for comparison between groups. Unpaired t-test (with Welch’s correction where appropriate), Mann-Whitney U, One-way ANOVA, or Brown-Forsythe and Welch ANOVA were used to compare experimental groups. Mixed-effects analysis was used to compared repeated time points between experimental groups.

## 3. Results

We first wanted to know whether non-cell-specific cellular processes of F11 cells were affected by radiation exposure, such as senescence, mitochondrial functions, oxidative stress, and cellular metabolism via the NAD/NADH^+^ pathway.

### 3.1. Superoxide dismutase (SOD) activity assay

Superoxide dismutase enzyme activity was compared between ionising radiation and sham exposure treatment. SOD activity was normalised to protein concentration of each sample. Radiation exposure did not change the SOD activity compared to the sham-exposed cells (unpaired t test: *P* = 0.3137; *t* = 1.067, *df* = 9, Figure 1A).

**Figure 1:**
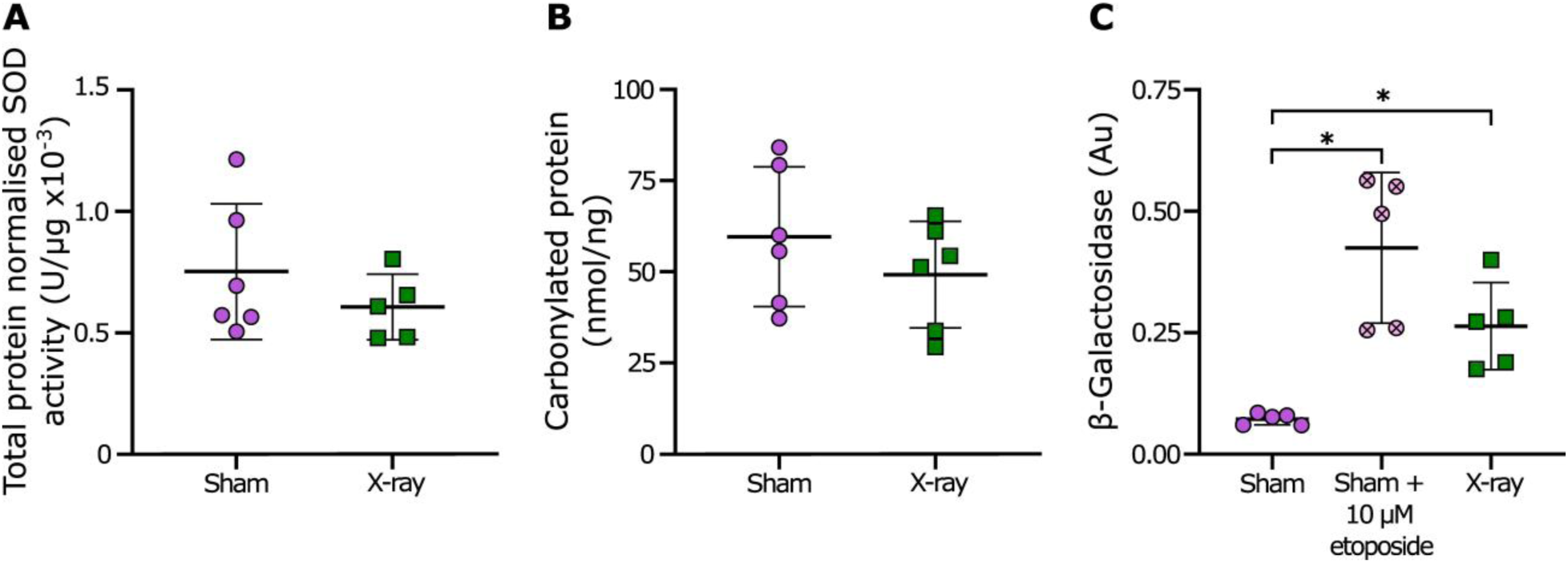
Oxidative stress and senescence response the day after 4 daily sham-or X-ray irradiations at 5 Gy. SOD activity normalized to total protein in U/µg x 10^-3^ (A) and carbonylated protein in nmol/ng (B) have been assessed as markers of oxidative stress. β-Galactosidase assay in arbitrary fluorescence units (Au) was used as a marker of senescence. Key: violet circle datapoints represent the sham irradiated experimental group, violet crossed circles represent the sham irradiated with the positive control etoposide (10 µM) experimental group, and green squares represent the X-ray irradiated experimental group. \**P* ≤ 0.05.

### 3.2. Carbonyl Assay

Protein oxidation was assessed by measuring the quantity of carbonyl groups. F11 cells exposed to ionising radiation did not show a change in quantity of carbonyl groups compared to sham-exposed cells (Unpaired t-test: *P* = 0.3163, *t* = 1.055, *df* = 10, Figure 1B).

### 3.3. Senescence

Senescence was measured by determining the β galactosidase activity. An increase of the β-galactosidase activity would represent an increase of senescence (Valieva *et al*., 2022).

There was a significant treatment difference (*P* = 0.0041; *F* (2, 6.474) = 14.48, Figure 1C) where the positive control (3 days exposure to 10 µM of etoposide) induced an increase of fluorescence intensity compared to the sham exposed group (*P* = 0.0181) but no difference to the X-ray exposed group (*P* = 0.2250). The X-ray exposed group had a significant increase of fluorescence intensity compared to the sham exposed group (*P* =0.0232).

### 3.4. Mitochondrial DNA copy

Exposure of F11 cells to X-rays resulted in a significant increase in relative mitochondrial DNA (mtDNA) copy number compared with sham-exposed cells (*P* = 0.0016; *t* = 4.285, *df* = 10, Figure 2A). The X-ray-exposed group showed a mean mtDNA relative copy number of 3021 ± 329.5 (n = 6), whereas the sham group showed 2296 ± 250.6 (n = 6).

**Figure 2:**
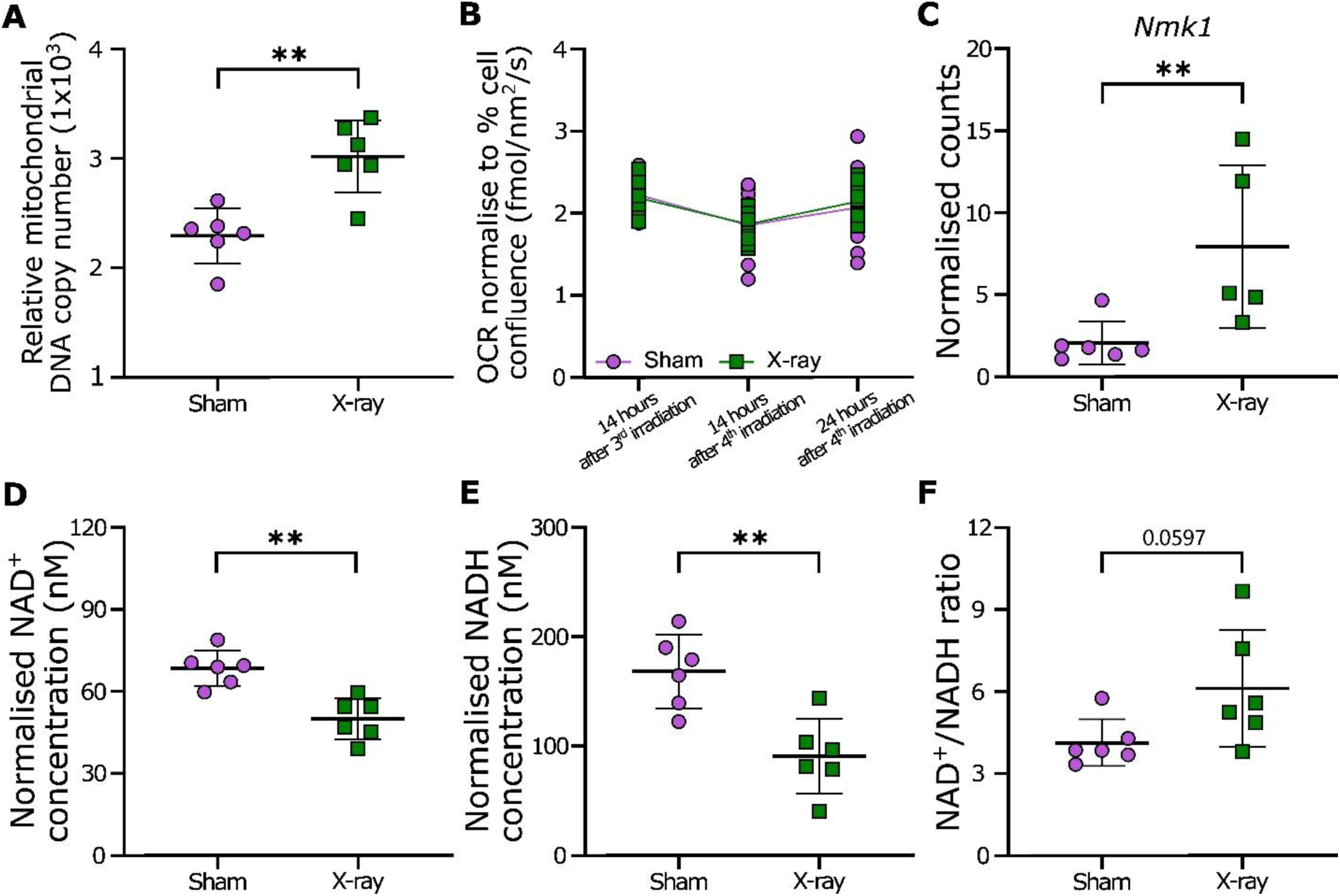
Mitochondrial functions and NAD^+^ salvage pathway comparison one day after 4 daily sham-or X-ray irradiations at 5 Gy. Relative mitochondrial DNA copy (A) and Oxygen consumption rate (OCR) expressed in fmol/nm² normalised to cell confluency (B) were used to assess aspects of mitochondrial functions. The OCR was measured for X-ray and sham groups 14h after the 3^rd^ and 4^th^ irradiation as well as 24h after the 4^th^ irradiation. *Nmrk1* gene expression (C), concentration of NAD^+^ (D) and NADH (E) expressed in nM normalised to cell confluency, and NAD^+^/NADH ratio (F) were used to assess the NAD^+^/NADH salvage pathway. Violet circles represent the sham irradiated treatment, and green squares represent the X-ray irradiated treatment. \*\**P* ≤ 0.01.

### 3.5. Oxygen consumption rate (OCR)

Sham-or X-ray-irradiated cells did not present differences in their oxygen consumption rate from the 3^rd^ radiation exposure to 24h after the last exposure (Figure 2B), only a time difference (Mixed-effects analysis: Time: P <0.0001, F (2, 44) = 30.33; Treatment: P = 0.8424, F (1, 30) = 0.04021; Time x Treatment: P = 0.5092, F (2, 44) = 0.6853).

### 3.6. NAD^+^/NADH pathway

*Nmrk1* gene expression was assessed as part of the biosynthesis of NAD+ via the salvage pathway (Xie *et al*., 2020). We observed increased *Nmrk1* expression in irradiated cells compared to sham (*P* = 0.0087, *U* = 1, Figure 2C).

Additionally, cellular concentrations of NAD^+^ and NADH normalised to cell confluency (normalised concentration expressed in nM) were calculated, and the ratio of NAD^+^/NADH was assessed. NAD^+^ (*P* < 0.0011, *t* = 4.542, *df* = 10, Figure 2D) and NADH (*P* = 0.0026, *t* = 3.974, *df* = 10, Figure 2E) were significantly reduced in irradiated cells while the ratio of NAD^+^/NADH (*P* = 0.0597, *t* = 2.123*, df* = 10, Figure 2F) was non-significantly increased yet with a tendency of increasing in irradiated cells compared to sham-exposed cells.

### 3.7. Radiation-induced gene expression changes in DRG neurons

There are 17 subpopulations of primary sensory DRG neurons: Cold, C-fibre low-threshold mechanoreceptors 1 (C-LTMR1), CLTMR2, Peptidergic fibres 1.1 (PEP1.1), PEP1.2, PEP2.1, PEP2.2, PEP2.3, Non-peptidergic fibres 1.1 (NP1.1), NP1.2, NP1.3, NP2.1, NP2.2, NP3, Aδ fibres, Aβ rapidly-adapting (RA) fibres, and proprio and Aβ slowly-adapting (SA) (Jung *et al*., 2023) associated with different sensory systems (mechanoreceptor, thermoceptor, proprioceptor, nociceptor, pruriceptor). Markers for subpopulations of DRG fibres are not specific to their subpopulation and differ between species. For instance, neurotrophic receptor tyrosine kinase 2 (NTRK2) is a marker of the Aδ-LTMR subpopulation but is also expressed in C-LTMR and cold subpopulations. Additionally, TAFA chemokine like family member 4 (TAFA4) shows a strong species difference in their specific fibre expression pattern (Jung *et al*., 2023). Similarly, the purinergic system involves both Aδ and C fibres with various mediators such as histamine, substance P, and transient receptor potential (TRP) channels (Misery *et al*., 2023). It is therefore difficult to pinpoint one gene and associated protein to an exclusive and specific DRG fibre and sensory system. However genetic markers exist to distinguish these subpopulations and their functional specialisations.

We started by analysing the expression of 6 genes present in DRG neurons, and the expression of the gene *P53* as a marker of radiation exposure using RT-qPCR. Neurotrophic receptor tyrosine kinase 1 (*Ntrk1*) is a marker of peptidergic nociceptors 2 (PEP2). *Ntrk2* is a marker of large diameter myelinated low-threshold mechanoreceptors A (A-LTMR fibres). Purinergic receptor P2Y1 (*P2ry1*) is present in C-fibre low-threshold mechanoreceptors (C-LTMR). Tachykinin precursor 1 (*Tac1*) is a marker of PEP but also cold thermoceptors. *Tafa4* is predominantly expressed in mouse, guinea pig and cynomolgus monkey C-LTMR and NP1 fibres while principally expressed in human Aδ, C-LTMR and cold fibres (Jung *et al*., 2023). Neuropilin 1 (*Nrp1*) is present in various DRG subpopulations (Gomez *et al*., 2023).

We observed that radiation exposure significantly increased the expression of the gene *Ntrk2* (*P* < 0.0001, *t* = 7.104, *df* = 9, Figure 3E) and *Tac1* (*P* = 0.0002, *t* = 6.067, *df* = 9, Figure 3F) 24 hours after the 4 daily doses of 5 Gy, but there were no effects on *Ntrk1* (*P* = 0.0573, *t* = 2.179, *df* = 9), *P2ry1* (*P* = 0.1255, Mann-Whitney U = 6), *Tafa4* (*P* = 0.0620, *t* = 2.168, *df* = 8), and *P53* (*P* = 0.2696, *t* = 1.176, *df* = 9, Figure 3D) when compared to sham-exposed cells (Table 2). Statistical analysis indicated a significant increase of *Nrp1* (*P* = 0.0174, *t* = 2.907, *df* = 9) in irradiated cells, however, the average fold change for the sham treated (0.9710 +/-0.05861) and the X-ray (1.113 +/-0.09853) were still within the variation of the RT-qPCR technique, and therefore not physiologically relevant (Table 2).

**Figure 3:**
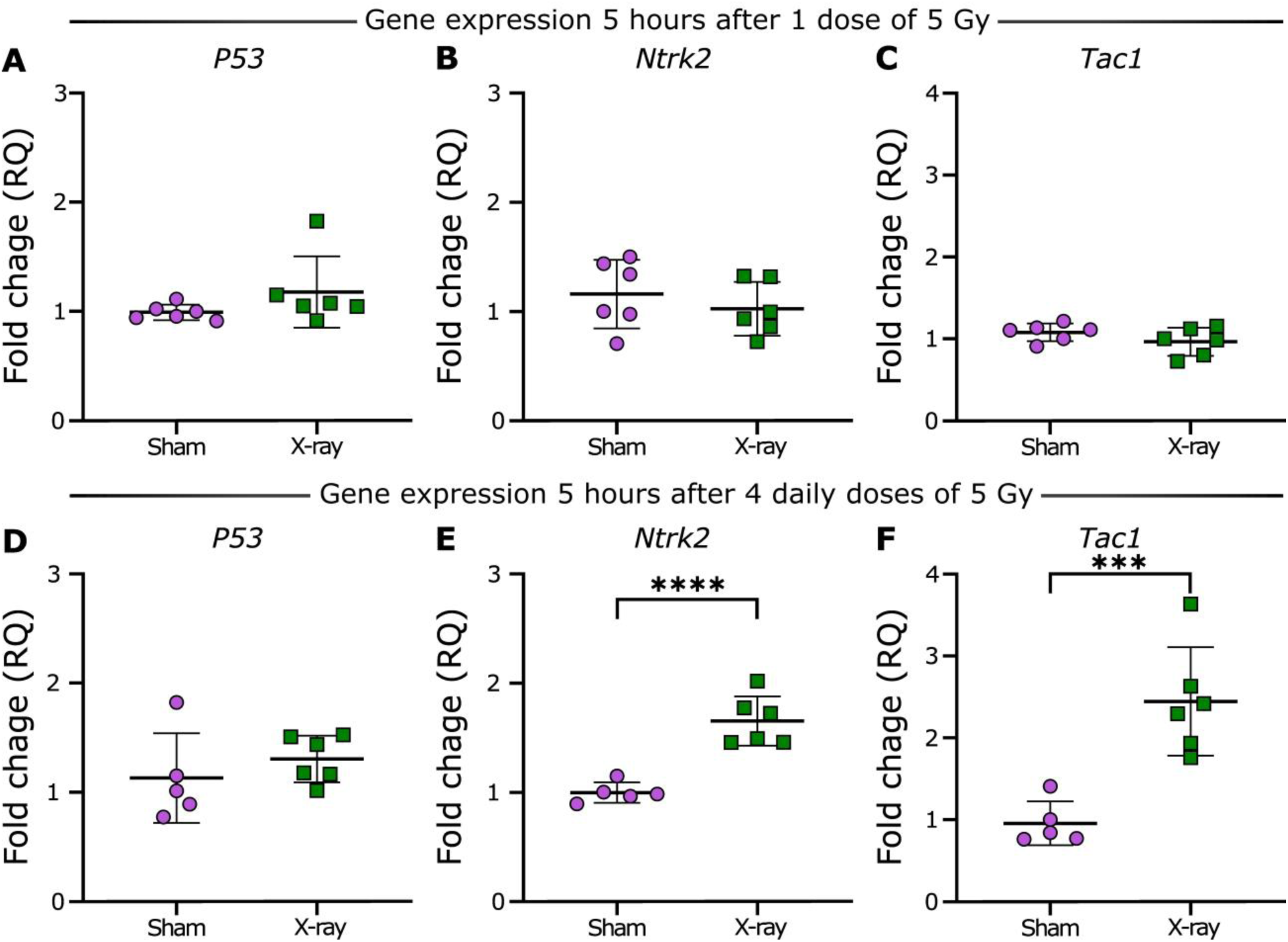
RT-qPCR gene expression analysis. Fold change (RQ) of *P53* (A), *Ntrk2* (B), and *Tac1* (C) is assessed 5 hours after 1 dose of sham or 5 Gy radiation exposure of F11 cells, and fold change (RQ) of *P53* (D), *Ntrk2* (E), and *Tac1* (F) was assessed 24 hours after 4 daily doses of sham or 5 Gy radiation exposure of F11 cells. The violet circles represent sham irradiated treatment, and green squares represent X-ray irradiated treatment. RQ = relative quantification, \*\*\**P* ≤ 0.001, ****P ≤ 0.0001.

**Figure 4:**
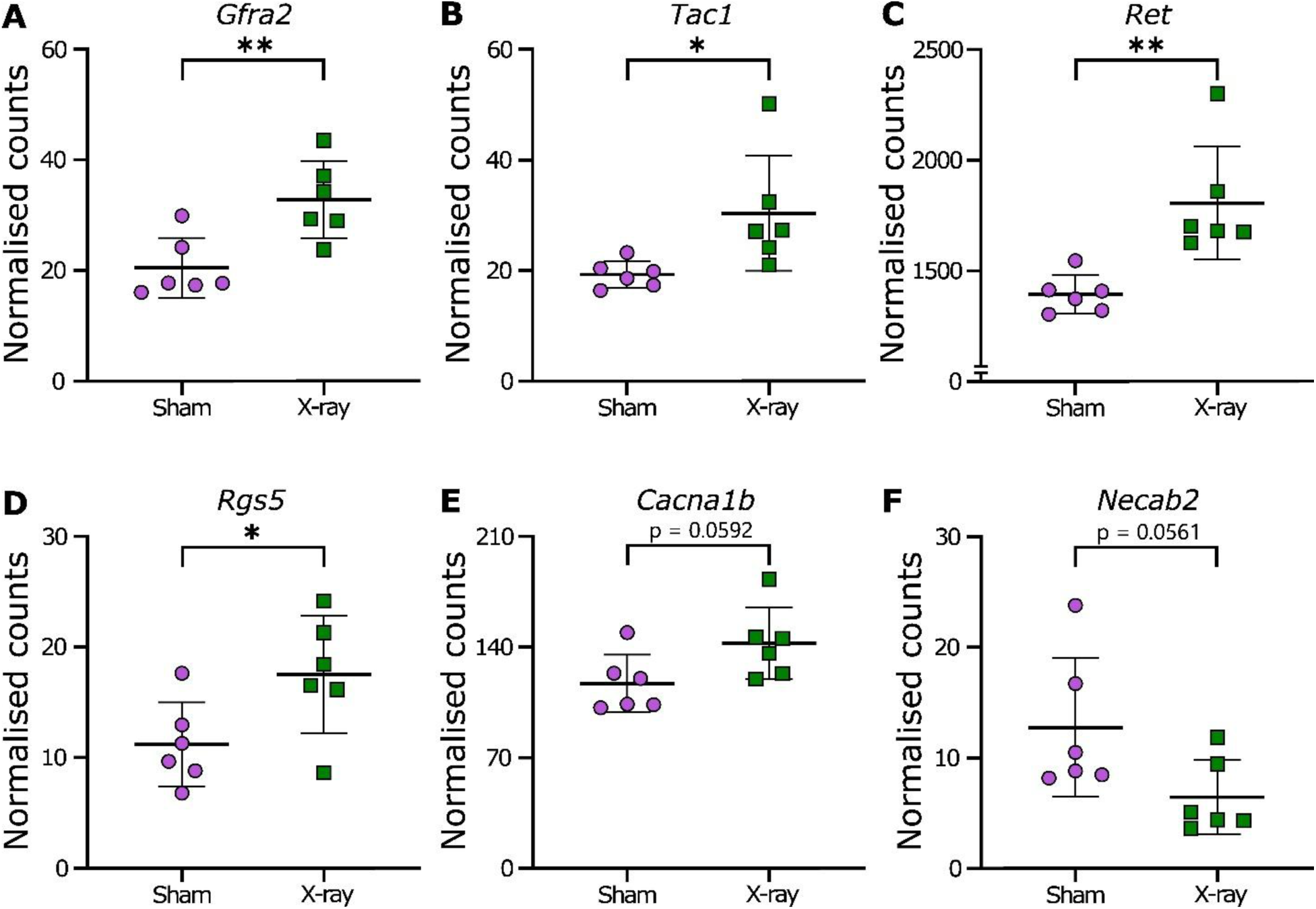
NanoString gene expression analysis. Changes in gene expression (normalised counts) of *Gfra2* (A), *Tac1* (B), *Ret* (C), *Rgs5* (D), *Cacna1b* (E), and *Necab2* (F) assessed 24 hours after 4 daily doses of sham or 5 Gy radiation exposure to F11 cells. Violet circles represent the sham irradiated treatment, and green squares represent the X-ray irradiated treatment. RQ = relative quantification, \**P* ≤ 0.05, \*\**P* ≤ 0.01.

**Table 2:** RT-qPCR gene expression analysis in sham-and radiation-exposed F11 cells. *Average fold change within the variation of the RT-qPCR technique, therefore not physiologically relevant.

| Gene | Effects | $P$, $t$, $df$ or $p$, $U$ | Mean control RNA expression (+/- standard deviation) | Mean X-ray RNA expression (+/- standard deviation) |
| --- | --- | --- | --- | --- |
| *Ntrk1* | Non-significant | $p = 0.0573$, $t = 1.179$, $df = 9$ | 2.131<br>($\pm 0.162$) | 1.854<br>($\pm 0.241$) |
| *Nrp1* | Statistically significant* | $p = 0.0174$, $t = 2.907$, $df = 9$ | 0.826<br>($\pm 0.088$) | 0.632<br>($\pm 0.125$) |
| *P2ry1* | Non-significant | $p = 0.1255$, $U = 6$ | 5.172<br>($\pm 0.525$) | 4.901<br>($\pm 0.167$) |
| *Tafa4* | Non-significant | $p = 0.062$, $t = 2.168$, $df = 8$ | 6.536<br>($\pm 0.518$) | 6.003<br>($\pm 0.181$) |

We also wanted to assess whether *P53*, *Tac1* and *Ntrk2* expression were affected 5h after a single exposure of 5 Gy. There was no significant difference between the sham control and irradiated cells for *P53* (*P* = 0.1320; *U* = 8, Figure 3A), *Ntrk2* (P = 0.3978, t = 0.8833, df = 10, Figure 3B), nor *Tac1* (P = 0.1902, t = 1.405, df = 10, Figure 3C) expression.

Following these data, we decided to assess whether X-ray exposure affected a broader transcriptional profile involved in DRG sensory neurons and discovered that radiation exposure affected six different genes out of 33 in F11 cells (Figure 5 and Supplement file 2).

### 3.8. Neuronal and sensory neuron markers

First, we observed no difference in two neuronal markers tubulin beta 3 class III (*Tubb3)* and synaptosome associated protein 25 (*Snap25)*. Additionally, no difference was noted in sodium voltage-gated channel alpha subunit 9 (*Scn9a)*, a marker for sensory neurons.

#### 3.8.1. Markers by pain modality

Among the genes associated with cold thermoception, there was no change between both experimental groups in the expression of forkhead box p2 (*Foxp2)*, nor retinoic acid receptor responder 1 (*Rarres1*). However, *Tac1* expressed in different neuronal subpopulation was increased in cells exposed to radiation. There was no change between experimental groups in the expression of acid-sensing ion channel (*Asic3*), which is involved in acidic pain (Deval *et al*., 2008). *Trpv1*, associated with noxious heat sensing, was not expressed in undifferentiated F11 cells. Moreover, *Trpm3*, also implicated in noxious heat transduction, exhibited no differences in expression among the experimental groups (Vangeel *et al*., 2020).

#### 3.8.2. Pruriceptive system

Within the panel of genes implicated in the pruriceptive system and itch signalling (Kahremany *et al*., 2020; Xing *et al*., 2020; Zhang *et al*., 2021; Mishra, 2022; Misery *et al*., 2023), no significant differences were detected between sham and irradiated F11 cells in the expression of histamine receptor h1 (*Hrh1*), glial cell derived neurotrophic factor family receptor alpha 1 (*Gfra1*), piezo type mechanosensitive ion channel component 1 (*Piezo1*), tyrosine hydroxylase (*Th*), transient receptor potential cation channel subfamily m member 2 (*Trpm2*), transient receptor potential cation channel subfamily v member 4 (*Trpv4*), contactin associated protein 2 (*Cntnap2*), solute carrier family 17 member 7 (*Slc17a7*), opioid receptor mu 1 *(Oprm1) nor* cannabinoid receptor 1 (*Cnr1*). In contrast, radiation exposure produced significant upregulation of regulator of G-protein signalling 5 (*Rgs5*) and *Tac1*.

#### 3.8.3. Markers by fibres

No difference was observed in *Cnr1,* mainly expressed in A-fibres (Hohmann & Herkenham, 1999; Bridges *et al*., 2003), nor *Oprm1* present in PEP1 fibres (Jung *et al*., 2023).

αδ and C-LTMRs share common gene markers. There was no change between experimental groups in the expression of sodium voltage-gated channel alpha subunit 5 (*Scn5a*), N-terminal EF-hand calcium binding protein 2 (*Necab2*)*, Rarres1* nor *P2ry1* expressed in Aδ and C-LTMRs. However, *Necab2* showed a non-statistically significant decrease in X-ray irradiated cells (*P* = 0.0561). More specific to C-LTMRs, there was no change in *Th* and diacylglycerol kinase eta (*Dgkh*), but regulators of *Rgs5* and *Gfra2* were increased in cells exposed to radiation.

Amongst the non-peptidergic fibre (NPs) markers, there were no changes between both experimental groups in the expression of *Gfra1*, nor in plexin C1 (*Plxnc1*). However, irradiated cells exhibited increased expression of *Gfra2* and *Ret*, genes expressed in populations of NPs (Jung *et al*., 2023), as well as in LTMR (Olson *et al*., 2016). Additionally, the expression of *Ntrk2*, present in Aβ RA-LTMR and Aδ-LTMRs (Olson *et al*., 2016), was also increased in irradiated cells. A-LTMR markers including *Necab2*, *Cntnap2*, *Slc17a7, Scn5a* and *Ntrk3* were not differentially expressed in both experimental groups, however *Necab2* was non-significantly decreased in the X-ray irradiated group (*P* = 0.0561). Expression of *Ret*, *Tac1*, and *Gfra2*, which are all markers of Aβ RA-LTMR (Luo *et al*., 2009), were increased in irradiated cells.

In addition, other genes implicated in pain signalling and nociception exhibited no change in expression. This included zinc finger homeobox-3 (*Zfhx3*), expressed in DRG neurons (Megat *et al*., 2019), which has been shown to be involved in heat and mechanical hypersensitivity in a preprint (Nolan *et al*., 2024); *Tafa4*, which has a species-dependent involvement in sensory DRG neurons; likewise for *Scn8a* (Jung *et al*., 2023). There was no change in calcium voltage-gated channel subunit alpha1 A (*Cacna1a*), but there was a non-significant (*P* = 0.0592) increase of the calcium voltage-gated channel subunit alpha1 B (*Cacna1b*) in irradiated cells.

## 4. Discussion

### 4.1. Cellular impact of radiation exposure

We first demonstrated that radiation induces senescence, a common outcome from radiotherapy that can ultimately lead to either cell death or cell survival (Li *et al*., 2018; Patel *et al*., 2020; Kim *et al*., 2023; Ibragimova *et al*., 2024). In this model, we observed senescence in F11 cells exposed to a cumulative dose of 20 Gy delivered over four consecutive days. This exposure dose represents the normal tissue dose constraints for stereotactic radiotherapy of the brainstem (Hanna *et al*., 2018). These findings therefore demonstrate that radiation at clinically relevant doses is sufficient to induce early cellular damage in this neuronal cell model.

Oxidative stress is a common cellular reaction to radiation (Zheng *et al*., 2023), yet we did not observe changes in the superoxide dismutase, nor in protein oxidation in cells exposed to radiation. We assessed both aspects of oxidative stress the day after the four-day irradiation regime. Therefore, it is possible that oxidative stress may have returned to a basal level due to cellular adaptation, despite the observed increase of mitochondrial DNA copy number, which is an early marker of cell response to oxidative stress (Lee *et al*., 2000). Further study is needed to confirm whether oxidative stress is present at earlier time points, especially when considering that SOD has radioprotective effects (Sonis, 2021; Xue *et al*., 2021), and reduces inflammatory (Bernardy *et al*., 2017) and oxaliplatin-induced pain (Guillaumot *et al*., 2019). Additionally, increased mitochondrial DNA copies protect from cell apoptosis (Cerritelli *et al*., 2003; Mei *et al*., 2015, 2020), thus the observed increase of mitochondrial DNA copy number could also be a cellular protection mechanism against radiation-induced apoptosis.

The dynamic response of P53 is linked to radioresistance and cell fates following radiation exposure. Transient increase of P53 activation is present in radioresistant tissues and cells (Stewart-Ornstein & Lahav, 2017; Stewart-Ornstein *et al*., 2021), and cells recovering from DNA damage (Purvis *et al*., 2012), while sustained P53 activity is present in radiosensitive cells and leads to senescence. However, this response is cell-and tissue-specific. In this investigation, *P53* expression was unaffected in irradiated groups, potentially indicating a radioprotective effect. In addition to radiation and DNA damage, p53 is also activated by oxidative stress (Liu & Xu, 2011; Shi *et al*., 2021). As we did not observe changes in oxidative stress, it would align with the lack of change to the *P53* transcript levels. Additional investigation is needed to determine whether the absence of *P53* modulation 24 hours after the last radiation dose reflects a form of radiosensitivity that favours protection from apoptosis, or whether this response is unique to the F11 cell line. Especially given that we did not observed changes at the acute single exposure of 5 Gy 5h after irradiation. This timepoint is justified by previous work, demonstrating a consistent *P53* response across various cell lines, but not including F11, 5 hours after radiation exposure despite p53 dynamics being cell specific (Stewart-Ornstein & Lahav, 2017).

Additionally, we observed an increase of *Nmrk1* in irradiated cells. NMRK1, also named NRK1, is a key factor in the synthesis of nicotinamide adenine dinucleotide (NAD^+^), which has neuroprotective properties against neurodegenerative diseases and axonal degeneration (Ratajczak *et al*., 2016; Braidy *et al*., 2019; Xie *et al*., 2020; Amjad *et al*., 2021; Covarrubias *et al*., 2021; Icso & Thompson, 2022), and diabetic neuropathy-induced DRG degeneration (Chandrasekaran *et al*., 2024). While proteins involved in the biosynthetic pathway of NAD^+^ can delay axonal degeneration, NMRK1 does not protect from axotomy (Sasaki *et al*., 2006). The radiation-induced upregulation of *Nmrk1* may help limit further neuronal loss, as NMRK1 contributes to NAD^+^ biosynthesis via the salvage pathway (Xie *et al*., 2020). This pathway is essential for maintaining cellular levels of NAD^+^ (Braidy *et al*., 2019), as depletion of NAD^+^ has been linked with aging, senescence, DNA damage repair dysfunction, metabolism and glycolysis dysfunction (Bai & Cantó, 2012; Amjad *et al*., 2021; Covarrubias *et al*., 2021), including cell death due to inability to process glucose as source of energy (Ying *et al*., 2005). Importantly, depletion of NAD^+^ is induced by nerve injury and further tissue damage (Metcalfe *et al*., 2023), moreover NAD^+^ is essential to cell survival (Covarrubias *et al*., 2021). In addition, ionising radiation has been shown to induce post-translational activation of nicotinamide phosphoribosyltransferase (NAMPT), another key enzyme of the salvage pathway, to restore NAD^+^ levels and support both DNA-damage repair and cell survival (Liao *et al*., 2022). These adaptive responses could account for the maintenance, with an increased tendency, of NAD^+^/NADH ratio observed in irradiated F11 cells, despite the reductions in total NAD^+^ and NADH levels in the F11 DRG neurons following the irradiation.

It is noteworthy that the NAD^+^ pathway is also involved in neuronal differentiation, especially via NTRK2 interaction (Neves *et al*., 2022). *Ntrk2* expression was also increased in the radiation-exposed F11 cells. At this stage, it would be speculative to hypothesise that the increase of *Nmrk1* and *Ntrk2* could be linked to DRG neuronal remodeling, but further experiments would be necessary to develop this avenue.

### 4.2. DRG gene expression changes due to radiation exposure

DRG neurons contain a complex bundle of specialised sensory fibres. These fibres vary in their degree of myelination, from unmyelinated C-fibres to large-diameter heavily myelinated A-fibres, and process diverse sensory modalities, including nociception, touch, thermoception, proprioception, and also pruriception (itch). Although genetic markers exist to distinguish these subpopulations and their functional specialisations, comprehensive characterisation remains challenging, especially given species-dependent differences in gene expression and sensory-neuron organisation. 17 transcriptionally distinct DRG sensory neuron subtypes have been identified (Jung *et al*., 2023). Amongst a selection of 82 genes expressed in various subtypes of DRG sensory neurons, only 35 were expressed in this F11 cell line. Whilst it is not possible to establish a full picture of radiation-induced gene expression changes in DRG sensory neurons, we were able to detect an increase in the expression of seven genes as a result of radiation exposure: *Ntrk2*, *Tac1*, *Nmrk1*, *Gfra2*, *Ret*, *Rgs5*, and *Nrp1*. Additionally, there was also a non-significant increase of *Cacna1b* and *Necab2* expression. We were also able to confirm that the increase of *Ntrk2* and *Tac1* gene expression was absent 5 hours after an acute exposure of 5 Gy, showing that the changes observed are likely due to long-term X-ray exposure.

Interestingly, with the exception of *Rgs5*, genes associated with DRG neurons influenced by radiation exposure are all expressed in Aβ LTMR (Luo *et al*., 2009; Olson *et al*., 2016; Gomez *et al*., 2023; Jung *et al*., 2023). Ret Proto-Oncogene (*Ret*) is a marker of Aβ RA-LTMR fibres. More precisely, early DRG neurons expressing *Ret*, *Gfa2* but not *TrkA* were Aβ RA-LTMR innervating Meissner’s corpuscles (Luo *et al*., 2009). Aβ LTMR are mechanoreceptors but with pro-nociceptive properties (Tashima *et al*., 2018; Gautam *et al*., 2024) involved in pain due to injury (Gangadharan *et al*., 2022) or chronic pain conditions (Israel *et al*., 2025). Activation of the Aβ LTMR in mouse skin has a pro-nociceptive effect in inflammatory and neuropathic pain models (Gautam *et al*., 2024). The authors also observed an electrophysiological shift of Aβ RA-LTMR fibres becoming like Aβ SA-LTMR fibres, and a reduction of the innervation of the Meissner’s corpuscles in mice with chronic inflammation (Gautam *et al*., 2024). Nerve injury also increases the expression of *Cacna1b* in DRG mechanoreceptors (Nieto-Rostro *et al*., 2023) and decreases the expression of *Necab2* in DRG neurons (Zhang *et al*., 2014), both observed as a non-statistical significant trends in our X-ray irradiated cells. Chronic pain can be induced by radiotherapy (Chua & Vig, 2023) and we have indicated that radiation induces changes in gene expression associated with Aβ LTMR fibres. Aβ LTMR fibres becoming involved in pain and nociception could be a potential mechanism in radiation-induced chronic pain that warrants further investigation.

### 4.3. Caveat

The F11 cell line is a hybrid cell line from mice neuroblastoma and rat embryonic DRG neurons. It is possible to differentiate F11 cells by serum deprivation for two weeks, however about half of the neuron population dies during the process (Hashemian *et al*., 2017; Pastori *et al*., 2019). Consistent with previous reports that serum withdrawal induces cell death in F11 cells (Linnik *et al*., 1993), neurons differentiated via serum deprivation exhibited compromised viability by the end of the radiation protocol. Cell lines do not always express all markers associated with their cell type, and it is particularly true of complex cell lines with various subpopulations like DRG neurons (Haberberger *et al*., 2020). Undifferentiated and differentiated F11 models are imperfect (Hashemian *et al*., 2017; Haberberger *et al*., 2020), but F11 cells provide an accessible way to assess radiation exposure effects *in vitro*, as a preliminary investigation of the nociception system.

Beta galactosidase activity is commonly used to assess senescence, however differentiated neuronal cells, such as DRG neurons, do not fulfil one of the senescence phenotypes associated with cell cycle arrest. Therefore, we should use precaution in assessing neurosenescence (Hudson *et al*., 2024). However, the use of an undifferentiated immortalised neuronal cell line offers the advantage of enabling the application of commonly used senescence biomarkers (Itahana *et al*., 2007).

## 5. Conclusion

DRG neuronal cells exposed to radiation doses equivalent to brainstem normal-tissue dose constraints used in stereotactic radiotherapy exhibited characteristic features of radiation-induced injury. Specifically, radiation induced senescence, changes in mitochondrial copy number, and changes in cell metabolism via the NAD^+^/NADH pathway, while leaving oxygen consumption rate, and oxidative stress levels unchanged. Furthermore, radiation exposure modulated the expression of a defined gene set enriched in Aβ LTMR, suggesting a potential sensitisation of this DRG subpopulation, which is known to contribute to pain during tissue injury and in chronic pain conditions.

## Authors’ contribution

WT: Investigation, Methodology, Writing – Review & Editing

SMG: Investigation, Visualization, Writing – Review & Editing

RaS: Investigation

CDL: Investigation

RiS: Investigation, Visualization, Writing – Review & Editing

MN: Investigation

NB: Conceptualization, Formal Analysis, Investigation, Methodology, Project Administration, Resources, Supervision, Validation, Writing – Original Draft Preparation, Writing – Review & Editing

## Acknowledgments

We thank Dr. John Byrne for their advice and discussion about the physics of radiotherapy. We also thank *brainstrust* and their participants for providing advice and comments on our research projects.

