## Supplementary File 2 for "Fractionated ionising radiation affects cellular functions, and gene expression associated to subpopulation of F11 dorsal root ganglia neurons without inducing oxidative stress"

Supplement file 2: NanoString gene expression analysis. Gene expression intensity is expressed in normalised counts.

| Gene | Effects | ***p*, *t*, *df*** or **p, *u*** | **Mean RNA expression intensity control (+/- standard deviation)** | **Mean RNA expression intensity x-ray (+/- standard deviation)** |
|---|---|---|---|---|
| ***Asic3*** | No difference | *P* = 0.1645, *t* = 1.513, *df* = 9 | 3.822 (± 1.168) | 2.728 (± 1.214) |
| ***Cacna1a*** | No difference | *P* = 0.1302, *t* = 1.649, *df* = 10 | 45.10 (± 7.126) | 54.19 (± 11.47) |
| ***Cacna1b*** | No difference | *P* = 0.0592, *t* = 2.129, *df* = 10 | 117.1 (± 18.32) | 142.5 (± 22.70) |
| ***Cnr1*** | No difference | *P* = 0.1949, *t* = 1.389, *df* = 10 | 440.5 (± 23.97) | 487.7 (± 79.71) |
| ***Cntnap2*** | No difference | *P* = 0.2449, *t* = 1.236, *df* = 10 | 125.3 (± 11.64) | 135.8 (± 17.27) |
| ***Dgkh*** | No difference | *P* = 0.7381, *t* = 0.3438, *df* = 10 | 15.34 (± 2.364) | 16.69 (± 9.322) |
| ***Foxp2*** | No difference | *P* = 0.4142, *t* = 0.8519, *df* = 10 | 11.29 (± 5.013) | 13.23 (± 2.425) |
| ***Gfra1*** | No difference | *P* = 0.6758, *t* = 0.4308, *df* = 10 | 17.37 (± 4.394) | 18.56 (± 5.146) |
| ***Gfra2*** | Increased in irradiated cells | *P* = 0.0067, *t* = 3.409, *df* = 10 | 20.47 (± 5.424) | 32.80 (± 7.005) |
| ***Hrh1*** | No difference | *P* = 0.2368, *t* = 1.259, *df* = 10 | 8.342 (± 3.603) | 11.92 (± 5.956) |
| ***Necab2*** | No difference | *P* = 0.0561, *t* = 2.160, *df* = 10 | 12.76 (± 6.284) | 6.470 (± 3.373) |
| ***Nmrk1*** | Increased in irradiated cells | *P* = 0.0087, *Mann-Whitney U = 1* | 2.062 (± 1.301) | 7.938 (± 4.951) |
| ***Ntrk3*** | No difference | *P* = 0.4726, *t* = 0.7464, *df* = 10 | 19.48 (± 6.256) | 22.09 (± 5.850) |
| ***Oprm1*** | No difference | *P* = 0.8182, *Mann-Whitney U = 16* | 38.10 (± 7.226) | 40.76 (± 9.786) |

| | | | | |
|---|---|---|---|---|
| ***P2ry1*** | No difference | *P* = 0.9057, *t* = 0.1215, *df* = 10 | 4.652 (± 3.616) | 4.870 (± 2.509) |
| ***Piezo1*** | No difference | *P* = 0.0973, *t* = 1.829, *df* = 10 | 37.20 (± 10.49) | 48.94 (± 11.71) |
| ***Plxnc1*** | No difference | *P* = 0.0708, *t* = 2.022, *df* = 10) | 104.2 (± 13.50) | 123.8 (± 19.43) |
| ***Rarres1*** | No difference | *P* = 0.4070, *t* = 0.8655, *df* = 10 | 5.415 (± 3.290) | 3.908 (± 2.712) |
| ***Ret*** | Increased in irradiated cells | *P* = 0.0022, *Mann-Whitney U = 0* | 1394 (± 86.54) | 1807 (± 254.4) |
| ***Rgs5*** | Increased in irradiated cells | *P* = 0.0386, *t* = 2.381, *df* = 10 | 11.22 (± 3.793) | 17.56 (± 5.307) |
| ***Scn5a*** | No difference | *P* = 0.1883, *Mann-Whitney U = 7.5* | 3.880 (± 1.627) | 1.888 (± 1.592) |
| ***Scn8a*** | No difference | *P* = 0.0782, *t* = 1.962, *df* = 10 | 7.637 (± 3.717) | 3.873 (± 2.875) |
| ***Scn9a*** | No difference | *P* = 0.2570, *t* = 1.202, *df* = 10 | 4.840 (± 1.847) | 7.088 (± 4.192) |
| ***Slc17a7*** | No difference | *P* = 0.7922, *Mann-Whitney U = 13* | 2.308 (± 1.349) | 2.742 (± 1.937) |
| ***Snap25*** | No difference | *P* = 0.208, *t* = 1.346, *df* = 10 | 542.6 (± 38.99) | 501.9 (± 62.91) |
| ***Tac1*** | Increased in cell irradiated | *P* = 0.0476, *t* = 2.533, *df* = 5.553 | 19.27 (± 2.450) | 30.33 (± 10.40) |
| ***Tafa4*** | No difference | *P* = 0.0554, *t* = 2.168, *df* = 10 | 7.748 (± 2.995) | 4.593 (± 1.935) |
| ***Th*** | No difference | *P* = 0.3095, *Mann-Whitney U = 11* | 80.42 (± 5.961) | 83.10 (± 8.560) |
| ***Trpm2*** | No difference | *P* = 0.3939, *Mann-Whitney U = 12* | 26.62 (± 5.820) | 23.01 (± 5.399) |
| ***Trpm3*** | No difference | *P* = 0.9372, *Mann-Whitney U = 17* | 6.167 (± 2.774) | 5.832 (± 3.768) |

| | | | | |
|---|---|---|---|---|
| ***Trpv4*** | No difference | *P* = 0.7644, *t* = 0.3080, *df* = 10 | 9.072 (± 1.857) | 8.520 (± 3.975) |
| ***Tubb3*** | No difference | *P* = 0.1718, *t* = 1,472, *df* = 10 | 827.6 (± 59.06) | 889.1 (± 83.52) |
| ***Zfhx3*** | No difference | *P* = 0.9696, *t* = 0.03913, *df* = 10 | 240.6 (± 24.02) | 241.5 (± 53.15) |
